# Atypical soluble guanylyl cyclases control brain size in *Drosophila*

**DOI:** 10.1101/2024.07.17.603894

**Authors:** Daniel Prieto, Boris Egger, Rafael Cantera

## Abstract

Hypoxia-induced proliferation of neural stem cells has a crucial role in brain development. In the brain of *Drosophila melanogaster*, the optic lobe exhibits progressive hypoxia during larval development. Here, we investigate an alternative oxygen-sensing mechanism within this brain compartment, distinct from the canonical hypoxia signaling pathway mediated by HIF. Using genetic tools, immunostaining, and confocal microscopy, we demonstrate that the loss of the atypical soluble guanylyl cyclase (asGC) subunit *Gyc88E*, or the ectopic expression of *Gyc89Db* in neural stem cells leads to increased optic lobe volume. We propose the existence of a link between cGMP signaling and neurogenesis in the developing brain.

## Description

Oxygen availability is a powerful driver of evolutionary novelty, and metazoans have developed sophisticated hypoxia sensing systems that are intricately related to the control of stem cell niches and development (Simon and Keith 2008; Rytkönen et al. 2011; Hammarlund et al. 2018). Increased atmospheric oxygen levels during evolution had profound effects on insects enabling them, for instance, to develop larger body size and the ability to fly (Stamati et al. 2011).

The atypical soluble guanylyl cyclase (asGC) subunits Gyc88E, Gyc89Da and Gyc89Db are expressed from embryo to adult stages in *Drosophila melanogaster* (Langlais et al. 2004; Morton et al. 2005). They are regulated by O_2_ levels and when activated by hypoxia generate cGMP, functioning as molecular O_2_ sensors (Vermehren et al. 2006) with faster responses than the canonical hypoxia pathway (de Lima et al. 2021).

The neural stem cells within the optic lobe (OL) of the *Drosophila* larval brain are hypoxic relative to the central brain (Misra et al. 2017), with different cell types experiencing varying degrees of hypoxia (Baccino-Calace et al. 2020). There is evidence that optic lobe progenitor cells might not exhibit a canonical hypoxia response (Baccino-Calace et al. 2020). *In situ* hybridization data (Langlais et al. 2004) suggests that asGC subunits are expressed within the OL, potentially linking O_2_ sensing to an alternative signaling pathway activated by hypoxia. Furthermore, transcriptomic data analysis (Yang et al. 2015), revealed that *Gyc89Db* is expressed in L3 neuroblasts. The combination of different asGC subunits might provide different sensitivity thresholds to hypoxia to *Drosophila* neurons (Lu et al. 2024).

Here, we used genetic tools to investigate the role of asGC in larval brain development. We explored asGC loss-of-function using a *Gyc88E*^*-/-*^ mutant and a *Gyc89Da*^*-/-*^*Db*^*-/-*^ double mutant. Additionally, we performed targeted gain-of-function with drivers specific to ectopically express *Gyc89Db* in neuroepithelial cells or overexpressing it in neuroblasts.

As the hypoxic OL remains neurogenic during larval life, we hypothesized that this hypoxia might activate asGC. Given that cGMP derived from nitric oxide signaling can activate mammalian neural stem cell proliferation (Santos et al. 2014), if hypoxia-driven cGMP signaling were to activate neural stem cell proliferation, loss of asGC function would abolish it. Contrary to our expectation, the *Gyc88E*^*-/-*^ loss-of-function mutant exhibited a 1.9-fold increase in brain hemisphere volume (Figure 1A). Interestingly, the double mutation *Gyc89Da*^*-/-*^*Db*^*-/-*^ had not the same effect. Notably, ectopic expression in NE or overexpression of *Gyc89Db* in NB lead to an increase in overall brain size (Figure 1A). The larval brain comprises two functionally and developmentally different compartments, the central brain (CB) which is formed mostly during embryonic stages and contains the neuronal synapses and the optic lobe (OL), which lack fully differentiated neurons with synapses and remains neurogenic throughout larval stages (Hartenstein et al. 2008; Baccino-Calace et al. 2020).

**Figure 1.**
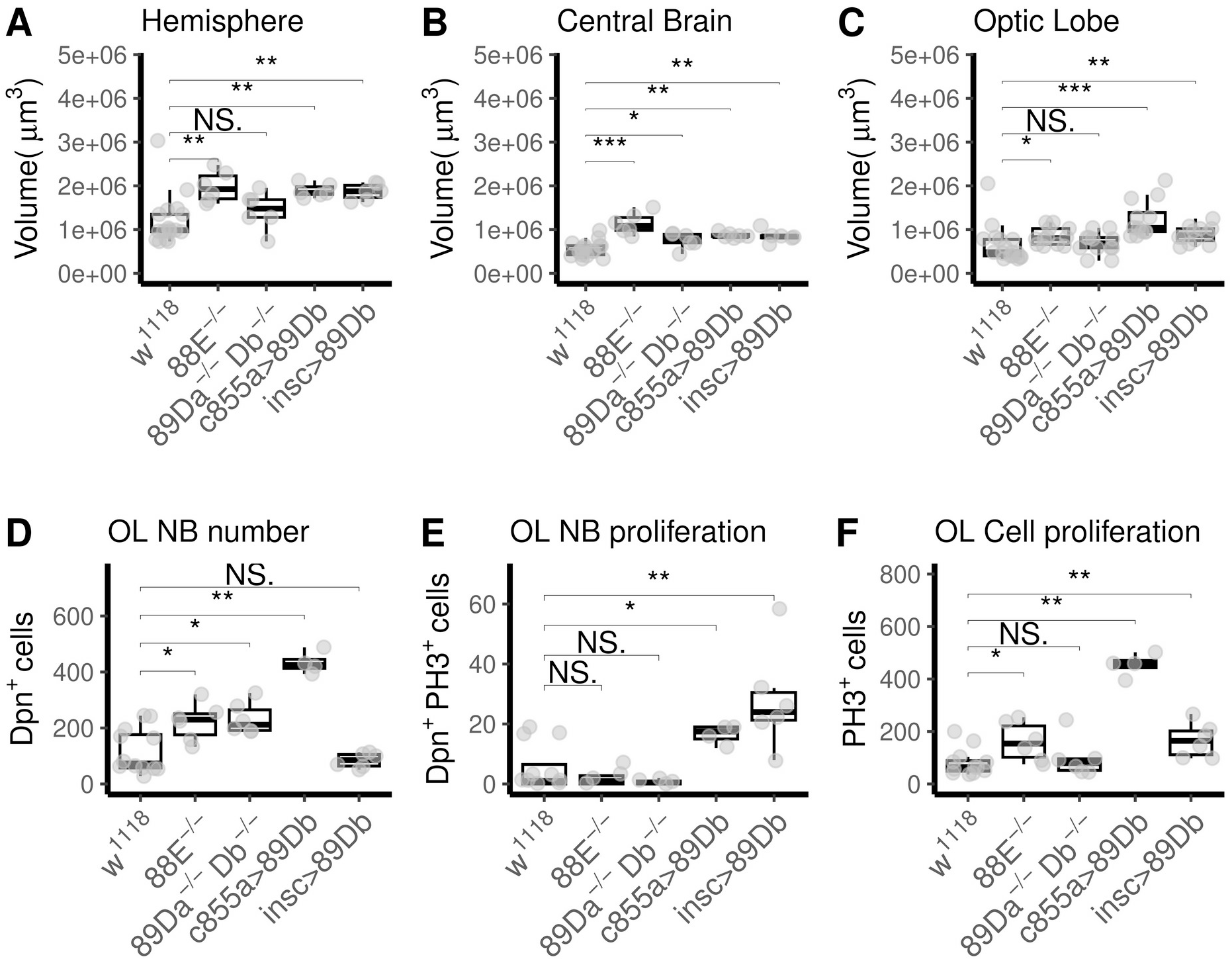
Effect of atypical soluble guanylyl cyclase loss- and gain-of-function in brain development of *Drosophila* larvae. **(A)** Volume of brain hemispheres at third larval instar. Loss-of-function *Gyc88E*^*-/-*^ mutant larvae (88E^-/-^) displayed increased brain hemisphere volume, (median±SD: 1.93e+06 ± 3.54e+05 μm^3^), whereas *Gyc89Da*^*-/-*^*Db*^*-/-*^ double mutant (89Da^-/-^Db^-/-^) did not (median±SD: 7.29e+05 ± 4.34e+05 μm^3^). Increased hemisphere volume was also observed in larvae ectopically expressing *Gyc89Db* in optic-lobe neuroepithelial cells under control of *GAL4*^*c855a*^ (c855a>89Db, median±SD: 1.84e+06 ± 1.53e+05 μm^3^) or overexpresing it in neuroblasts under control of *insc-Gal4* (insc>89Db, median±SD: 1.88e+06 ± 1.85e+05 μm^3^), compared to *w*^*1118*^ (median±SD: 9.86e+05 ± 5.70e+05 μm^3^). **(B)** Volume of central brain. Loss-of-function mutant larvae displayed increased central brain volume, *Gyc88E*^*-/-*^ (median±SD: 1.08e+06 ± 2.46e+05 μm^3^), *Gyc89Da*^*-/-*^*Db*^*-/-*^ (median±SD: 8.53e+05 ± 1.81e+05 μm^3^) as did ectopic expression lines driving *Gyc89Db* in optic-lobe neuroepithelial cells (median±SD: 8.53e+05 ± 6.18e+04 μm^3^). Overexpression in neuroblasts did not alter volume occupied by neuroblasts (median±SD: 8.41e+05 ± 1.37e+05 μm^3^), compared to *w*^*1118*^ (median±SD: 5.05e+05 ± 1.70e+05 μm^3^). **(C)** Volume of the optic lobe (OL). *Gyc88E*^*-/-*^ mutant larvae did not show increased OL volume (median±SD: 8.04e+05 ± 2.01e+05 μm^3^), nor did *Gyc89Da*^*-/-*^*Db*^*-/-*^ larvae (median±SD: 6.84e+05 ± 2.46e+05 μm^3^). Larvae ectopically expressing *Gyc89Db* in neuroepithelial cells showed increased OL volume (median±SD: 1.04e+06 ± 4.21e+05 μm^3^) whereas those overexpressing *Gyc89Db* in neuroblasts did not (median±SD: 8.72e+05 ± 1.95e+05 μm^3^) when compared to *w*^*1118*^ (median±SD: 4.81e+05 ± 4.23e+05). **(D)** Assessment of neuroblast number using anti-Deadpan immunostaining. Optic lobe neuroblasts were counted in the brain of larvae from each experimental group. The number of neuroblasts was increased in *Gyc88E*^*-/-*^ (median±SD: 230.00 ± 67.56) and in *Gyc89Da*^*-/-*^*Db*^*-/-*^ mutant larvae (median±SD: 211.50 ± 56.10). Larvae expressing *Gyc89Db* in neuroepithelial cells exhibited a 5.8-fold increase in their neuroblast number (median±SD: 425.50 ± 39.99). Overexpression in neuroblasts did not alter their number (median±SD: 85.00 ± 26.06) compared to *w*^*1118*^ (median±SD: 73.00 ± 79.18)larvae. **(E)** Quantification of proliferating optic lobe neuroblasts, identified by double staining for anti-Deadpan and anti-PH3. The number of proliferating neuroblasts was low, without statistically significant differences in *Gyc88*E^-/-^ (median±SD: 1.00 ± 2.76) or *Gyc89Da*^*-/-*^*Db*^*-/-*^ double mutant larvae (median±SD: 5.00e-01 ± 8.16e-01) compared to *w*^*1118*^ (median±SD: 1.00 ± 7.63). Brains expressing *Gyc89Db* under the neuroepithelial driver *GAL4*^*c855a*^ (median±SD: 17.5 ± 3.32) showed a 2.5-fold increase in the number of mitotically active neuroblasts, while those expressing Gyc89Db under the neuroblast driver *insc*-*gal4* showed a 3.4-fold increase (median±SD: 24.00 ± 16.76). **(F)** Quantification of total proliferating cells, measured by anti-PH3 immunostaining. *Gyc88E*^*-/-*^ larvae showed increased numbers of total proliferating cells (1.4-fold, median±SD: 154.00 ± 74.42) whereas *Gyc89Da*^*-/-*^*Db*^*-/-*^ did not (median±SD: 79.50 ± 74.46). Larvae expressing *Gyc89Db* in neuroepithelial cells under control of the *GAL4*^*c855a*^ driver exhibited a 4.3-fold increase in the number of mitotic cells (median±SD: 459.50 ± 44.14), whilst larvae expressing *Gyc89Db* under the neuroblast driver *insc-gal4* showed a 1.5-fold increase (median±SD: 164.50 ± 65.58) compared to *w*^*1118*^ larvae (median±SD: 64.50 ± 51.07). Mann-Whitney U test, * p<0.05, ** p<0.01, *** p<0.001. Mutant, ectopic expression, and overexpression N=6, *w*^*1118*^ N=12.

When we measured and compared CB and OL volumes separately, we observed that the macrocephalous phenotype observed in *Gyc88E*^*-/-*^ mutants can be explained by a specific effect on the CB. *Gyc88E*^*-/-*^ loss-of-function mutants 2.1-fold increase in CB volume (Figure 1B), while the OL volume remains unaffected (Figure 1C). The double mutation *Gyc89Da*^*-/-*^*Db*^*-/-*^ also elicited an increase in CB volume but limited to a lesser extent (1.7-fold, Figure 1B).

Ectopic expression of *Gyc89Db* under the control of the OL-specific neuroepithelial driver *GAL4*^*c855a*^ (Egger et al. 2007) resulted in increased volume of both CB (Figure 1B) and OL (Figure 1C). Overexpression under the pan-neuroblast driver *inscuteable*-GAL4 reproduced the macrocephalous phenotype only in the CB (Fig. 1B). Provided the OL is the major neurogenic region within the larval brain, experiencing a dramatic volume increase between 24-72 hs after larval hatching (Baccino-Calace et al. 2020), we reasoned that, in the larva, an eventual increase in proliferation would only be observable in this hypoxic brain region.

Aiming to further investigate the cellular basis of the macrocephalous phenotype, we assessed neuroblast number and proliferation within the OL. asGC loss-of-function mutants and ectopic expression of *Gyc89Db* in the OL neuroepithelium increased neuroblast number in the optic lobe, while overexpression in neuroblasts did not (Figure 1D).

Despite the increased OL volume, no differences in the number of proliferating neuroblasts were observed in the loss-of-function mutants (Figure 1E), whereas *Gyc89Db* ectopic expression in neuroepithelial cells or overexpression in neuroblasts increased the number of mitotically active neuroblasts by 2.5-fold and 3.4-fold, respectively (Figure 1E).

Furthermore, larvae with *Gyc88E*^*-/*-^ loss of function or ectopic expression of *Gyc89Db* in the neuroepithelium or overexpression in neuroblasts show increased total cell number in mitosis in the optic lobe (Figure 1F), suggesting a link between oxygen-dependent cGMP signaling and the control of proliferation in neural progenitor cells.

Although our studies were performed using well characterized mutant lines (Vermehren-Schmaedick et al. 2010), we cannot rule out that background mutations might also be affecting our results. Future studies might also consider assaying these mutations in trans to a chromosomal deficiency.

Previous research has shown that *Gyc88E* is active in the absence of additional subunits, while *Gyc89Da* and *Gyc89Db* enhanced the activity of *Gyc88E* when co-expressed, suggesting that these enzymes acted likely as heterodimers (Morton and Vermehren 2007; Morton 2011). It is important to mention that *Gyc89Db* is expressed in the brain during larval development (Langlais et al. 2004) and produces cyclic GMP in response to low oxygen *in vitro* (Morton 2004; Vermehren et al. 2006). Thus, asGCs may fine tune the cellular response promoting proliferation through hypoxia-driven cGMP signaling.

## Methods

### Fly strains

Flies were raised on cornmeal medium at constant temperature 25°C and under 12:12 h light:darkness cycles as previously described (Ferreiro et al. 2018). Loss-of-function stocks *Gyc88E*^*-/-*^ (Z3-1083, from the Seattle tilling project) and the double mutant *Gyc89Da*^*-/-*^*Db*^*-/-*^ (Vermehren-Schmaedick et al. 2010) (BDSC #93108) were a kind gift of David Morton (Oregon Health & Science University, Oregon, USA). The following driver and responder lines were used: *c855a-GAL4* (BDSC stock #6990), *insc-GAL4* (BDSC stock #8751), *UAS-gyc89Db* (Vermehren-Schmaedick et al. 2010) (BDSC stock #93116) was a kind gift from David Morton (Oregon Health & Science University, Oregon, USA). *w*^*1118*^ was used as control as mutant and transgenic lines were built on a *white* background.

### Immunostaining and image acquisition

All experiments were performed on wandering third-instar larvae kept in normoxia conditions. Brains were dissected, fixed and immunostained as previously described (Baccino-Calace et al. 2020). Primary antibodies included a monoclonal mouse anti-Discs large antibody to outline the cells and allow for precise identification of the OL boundaries (1:20, DSHB #4F3, Developmental Studies Hybridoma Bank (DSHB), Iowa, USA), guinea pig anti-Deadpan to identify neuroblasts (1:2500, kind gift from Jürgen Knoblich, Eroglu et al. 2014) and rabbit anti-phosphorylated H3 histone antibody as a mitotic marker (anti-PH3, 1:200, Cell Signaling Technologies #9713). Fluorescent conjugated secondary antibodies Alexa488, Alexa568 and Cy5 were used (Thermo-Fisher). DNA was stained with 1 μg/ml Hoechst 33342 (Thermo-Fisher). Images were acquired with a Zeiss LSM800 Airyscan confocal microscope and processed with FIJI (Schindelin et al. 2012). Semi-automated OL segmentation was performed with TrakEM2 (Cardona et al. 2012). Analyses and illustrations were made using R version 4.4.0 on RStudio version 2023.06.01.

## Reagents

Stocks: Stocks obtained from the Bloomington Drosophila Stock Center (BDSC) or kindly provided by colleagues as stated under the Methods section.

*GAL4*^*c855*^ (*w[1118]; P{w[+mW*.*hs]=GawB}C855a*)

*insc-GAL4* (*w[*]; P{w[+mW*.*hs]=GawB}insc[Mz1407]*)

*UAS-gyc89Db* (*w[*]; P{w[+mC]=UAS-Gyc89Db*.*V}2*)

*Gyc88E*^*-/-*^ (point mutation V474M)

*Gyc89Da*^*-/-*^*Db*^*-/-*^ (w[*]; PBac{w[+mC]=RB}Gyc89Da[e01821]

Mi{GFP[E.3xP3]=ET1}Gyc89Db[MB03197])

## Acknowledgements

The authors would like to thank Dr. Maria Jose Ferreiro for her continued valuable help with fly care and fruitful discussions, and the Developmental Studies Hybridoma Bank for antibodies. Stocks obtained from the Bloomington Drosophila Stock Center (NIH P40OD018537) were used in this study.

## Funding

DP was a post-doctoral researcher at the Instituto de Investigaciones Biológicas Clemente Estable, Ministerio de Educación y Cultura, Uruguay. Grants FVF-2019-05 (Dirección Nacional de Innovación, Ciencia y Tecnología) awarded to DP and FCE_1_2019_1_156160, awarded to RC and support from Agencia Nacional de Investigación e Innovación to DP and RC are acknowledged.

## Notes

### Competing Interest Statement

The authors have declared no competing interest.

### Summary of Updates

-Abstract updated to correct truncated sentence. -Materials and Methods and Reagents section: Gyc88E-/- fly stock statement revised. -Nomenclature of the double mutant revised to clarify its 'double' status. -Figure 1 revised to unify y-axis scale in panels A,B and C.

